# Network dynamics of Broca’s area during word selection

**DOI:** 10.1101/478461

**Authors:** Christopher R Conner, Cihan M Kadipasaoglu, Harel Z Shouval, Gregory Hickok, Nitin Tandon

## Abstract

Current models of word-production in Broca’s area posit that sequential and staggered semantic, lexical, phonological and articulatory processes precede articulation. Using millisecond-resolution intra-cranial recordings, we evaluated spatiotemporal dynamics and high frequency functional interconnectivity between ventro-lateral prefrontal regions during single-word production. Through the systematic variation of retrieval, selection, and phonological loads, we identified specific activation profiles and functional coupling patterns between these regions that fit within current psycholinguistic theories of word production. However, network interactions underpinning these processes activate in parallel (not sequentially), while the processes themselves are indexed by specific changes in network state. We found evidence that suggests that pars orbitalis is coupled with pars triangularis during lexical retrieval, while lexical selection in Broca’s area is terminated via coupled activity with M1 at articulation onset. Taken together, this work reveals that speech production relies on very specific inter-regional couplings in rapid sequence in the language dominant hemisphere.

## Introduction

The rapid and precise production of words is critical to fluent language. This process is thought to rely crucially on interactions between the inferior frontal gyrus (IFG) and the primary motor cortex. In the 150 years since the description of this region by Paul Broca[1], the functional anatomy of the left IFG has been explored in great detail using anatomical methods, lesion analysis, and functional neuroimaging[2-4]. Yet, language processes occur on a spatio-temporal scale that is finer than these methods can provide[5]. Specifically, the neural mechanisms by which Broca’s area enables the rapid word selection and production that underpins human language are unknown. A number of linguistic models[6-16] propose that word production sequentially involves lexical retrieval (the activation of several possible relevant responses), response selection (the narrowing of choices to the most appropriate one), and finally, phonological encoding (the selection and sequencing of speech sounds)[17]. These models imply that the time scale of these constituent processes is quite slow, and that semantico-lexical, phonological and articulatory processes occur sequentially, with hand-off occurring from the end of one process, to enable the next[17, 18]. However, recent work, suggests that very early (within 200 ms) after input, acoustic structure and semantics are already being processed[13, 19, 20]. Additionally, functional imaging techniques have also been used to ascribe this stepwise ontology of word production to distinct sub-regions of Broca’s area, and can parsimoniously be interpreted as implying that linguistic process reside in a distinct cortical region[17, 18, 21-23].

Over the last decade, invasive recordings of human cortex have led to novel insights into language mechanisms[6, 24-27]. However, these studies have evaluated the spatial and temporal characteristics of individual regions in isolation, with limited, if any, analyses of the network behavior likely underpinning them[6]. Here, we seek to evaluate how linguistic processes are effected by networks connecting sub-regions involved in these processes, and whether changes in network state index the transition from one constituent process to the next.

To derive this network-based description of dynamics within the IFG during word production, we collected data in a large cohort (n=27) of patients with experimental conditions that varied retrieval, selection, and phonological loads. We specifically evaluated interactions between sub-regions of the IFG and motor cortex in the sub-central gyrus (sCG) during the intervals at which constituent linguistic processes might occur. Intermediary states identified in the functional network connecting the components of the IFG during object naming should form an empiric basis to refine spreading activation models[28-30], and might lead to the modification of existent theories that propose that (i) individual components of Broca’s area make distinct and strictly separable contributions to lexical retrieval and selection processes[6-9, 31, 32], and (ii) that processing occurs in a fairly strict rostro-caudal progression [10, 33] – namely that anterior pre-frontal regions unidirectionally exert control over posterior pre-motor and motor regions [11, 12].

## Materials and methods

Intracranial electrophysiological data were collected from 27 patients (17 female, mean age 34±12 years) undergoing subdural electrode (SDE) implants for localizing seizure onset sites (Table 1). Seventeen patients were implanted over the left hemisphere, five underwent electrode implants bilaterally and five over the right hemisphere, following procedures described previously [34]. All experimental procedures were reviewed and approved by the Committee for the Protection of Human Subjects (CPHS) of the University of Texas Houston Medical School (IRB #HSC-MS-06-0385), and written informed consent was obtained from all subjects. In all cases, the left hemisphere was dominant for language. This was demonstrated in 18 patients by an intra-carotid injection of sodium amytal (the Wada procedure)[35] for lateralization of language function. 24 patients underwent language mapping using fMRI techniques and all were found to be left-hemisphere dominant. Nineteen underwent language localization using cortical stimulation mapping, resulting in language production deficits in 17. The remaining two subjects were only mapped over the right hemisphere. In these subjects, no deficit was produced, and they were therefore classified as left-hemisphere dominant.

**Table 1.** Individual characteristics of the 27 patients enrolled in the study. In 17 subjects, electrode implantation was solely over the left (L) hemisphere; in 5 subjects, it was over the right (R), and in 5 subjects, SDEs were placed bilaterally (B). Laterality indices (LI) were computed using fMRI during the same language tasks (n=23). LI >0 implies left hemisphere dominance for language. In 18 subjects, the intra-carotid amytal test (IAT) was used to determine language laterality. Cortical stimulation mapping (CSM) was performed in 19 subjects, of which, 17 mapped over the left hemisphere and two over the right (nos. 19 and 24). In all tested subjects, either deficits were produced on the left (L), or they were not observed on the right (Neg). The Edinburgh Handedness Inventory revealed that 25 subjects were right-hand dominant, and 2 were left-hand dominant.

| Pt Num | Sex | Age | Site | LI | IAT | CSM | Hand | IQ |
| --- | --- | --- | --- | --- | --- | --- | --- | --- |
| 1 | F | 37 | L | 0.56 | L | L | R | 89 |
| 2 | M | 21 | L | 0.40 | L | L | R | 97 |
| 3 | M | 39 | L | 0.52 | n/a | L | R | 100 |
| 4 | F | 38 | L | 0.27 | L | L | R | 96 |
| 5 | M | 17 | L | 0.43 | L | L | L | 67 |
| 6 | F | 30 | L | 0.56 | L | L | R | 107 |
| 7 | F | 20 | L | 0.72 | n/a | L | R | 103 |
| 8 | F | 30 | L | 0.82 | L | L | R | 100 |
| 9 | F | 20 | L | 0.28 | n/a | L | R | 97 |
| 10 | F | 42 | L | 0.80 | L | L | R | 107 |
| 11 | F | 28 | L | 0.77 | L | L | R | 97 |
| 12 | F | 51 | L | 0.19 | L | L | R | 92 |
| 13 | M | 42 | L | 1.00 | L | L | R | 109 |
| 14 | M | 23 | L | 0.89 | L | L | R | 91 |
| 15 | F | 29 | L | 0.46 | L | n/a | R | 84 |
| 16 | F | 45 | L | n/a | n/a | L | R | 93 |
| 17 | F | 21 | L | n/a | n/a | L | R | 78 |
| 18 | M | 30 | R | 0.44 | L | n/a | R | 90 |
| 19 | M | 27 | R | 0.78 | n/a | Neg | R | 112 |
| 20 | M | 40 | R | 0.84 | L | n/a | R | 94 |
| 21 | F | 51 | R | 0.91 | L | n/a | R | 74 |
| 22 | F | 28 | R | 0.30 | n/a | n/a | L | 107 |
| 23 | F | 62 | B | 0.33 | L | n/a | R | 93 |
| 24 | M | 56 | B | 0.79 | n/a | Neg | R | 116 |
| 25 | M | 24 | B | n/a | L | n/a | R | 105 |
| 26 | F | 30 | B | 0.24 | n/a | n/a | R | 120 |
| 27 | F | 31 | B | n/a | L | L | R | 124 |

## Language tasks

Data were collected as each patient performed two language tasks: naming of visually presented common objects[36] and scrambled images (scrambled from the collection of object stimuli) (Fig 1). A large stimulus pool of object images (Table 2) was used to induce variation in word frequency[37], selection load (varying number of possible correct responses[36]), and lexical class of the word (object name vs. scrambled images). For object naming, they were instructed to respond with a single word (Table 2). In response to the scrambled images, subjects were asked to articulate, “scrambled”.

**Fig 1.**
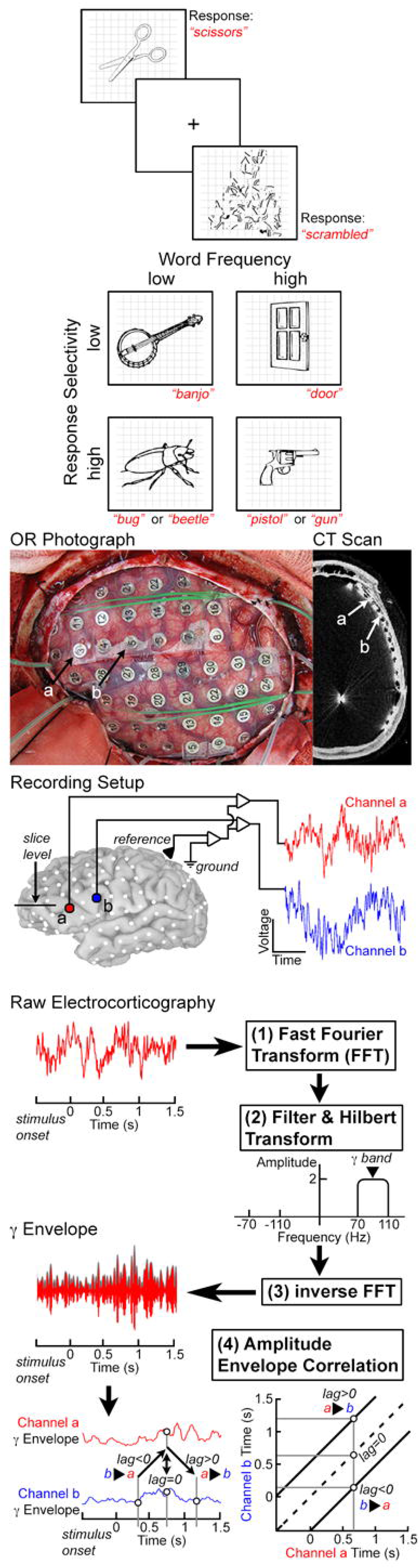
Experimental design. Two different classes of stimuli were used for visually cued generation of words: objects, and the control stimuli (scrambled versions of the target stimuli). Images were presented randomly, and subjects were asked to respond by saying, “*scrambled*” for control stimuli. Object stimuli were also ranked based on word frequency and selectivity (number of possible correct responses). Participants (n=27) were implanted with sub-dural electrodes (SDEs), and electro-corticographic (ECoG) data were recorded during task performance at 1-2 kHz. ECoG data were fast Fourier transformed (FFT) and filtered in the high gamma band (70-110 Hz). A Hilbert transform was applied, and data were then transformed back using an inverse FFT. This process generates an amplitude envelope of the desired frequency band (here the gamma band). Functional connectivity was assessed by correlating these envelopes of activity between two different channels – the amplitude envelope correlation (AEC). The amplitude envelopes across trials on one channel are correlated with a second channel to estimate the connectivity. Activity between two channels can be correlated at various lags to estimate the directionality of information flow. The dotted line on each graph represents a lag of 0 ms, while connectivity above or below this line is lagged from one region to the next (activity in region A affects region B at a later time point). This allows for the assessment of unidirectional connectivity from A to B and from B to A, along with bi-directional connectivity.

**Table 2.** List of all stimuli used in the analysis. Original images are available in Snodgrass, Joan G. and Vanderwart, Mary. "A standardized set of 260 pictures: Norms for name agreement, image agreement, familiarity, and visual complexity.” Journal of Experimental Psychology: Human Learning and Memory, Vol 6(2), Mar 1980, 174-215.http://dx.doi.org/10.1037/0278-7393.6.2.174

| Object Stimuli |  |  |  |  |  |  |
| --- | --- | --- | --- | --- | --- | --- |
| Airplane | Bowl | Cup | Globe | Mitten | Rhino | Table |
| Alligator | Box | Deer | Glove | Moon | Ring | Teapot |
| Anchor | Bread | Dog | Grapes | Mouse | Rocking-chair | Tennis racket |
| Ant | Broom | Doll | Grasshopper | Mushroom | Rooster | Thumb |
| Apple | Brush | Donkey | Guitar | Nail | Rose | Tie |
| Arm | Bus | Door | Gun | Necklace | Sailboat | Tiger |
| Artichoke | Butterfly | Doorknob | Hamburger | Needle | Sandwich | Toaster |
| Ashtray | Button | Dress | Hammer | Nose | Saw | Tomato |
| Asparagus | Cake | Drum | Hand | Nut | Scissors | Toothbrush |
| Axe | Camel | Duck | Hanger | Onion | Screw | Traffic light |
| Balloon | Candle | Eagle | Hat | Owl | Screwdriver | Train engine |
| Banjo | Cannon | Ear | Helicopter | Paintbrush | Seal | Tree |
| Banana | Cap | Elephant | Helmet | Pants | Shaker | Truck |
| Barn | Carrot | Envelope | Horse | Paperclip | Shell | Trumpet |
| Barrel | Cat | Eye | House | Peach | Shirt | Turtle |
| Baseball-bat | Celery | Fence | Hummingbird | Peanut | Shoe | Umbrella |
| Basket | Chain | Finger | Iron | Pear | Skate | Vase |
| Bat | Chair | Fish | Iron-board | Pelican | Skirt | Vest |
| Bear | Cherry | Flag | Jacket | Pen | Snowman | Violin |
| Bed | Chicken | Flower | Kangaroo | Pencil | Sock | Wagon |
| Bee | Cigar | Fly | Key | Penguin | Spider | Watch |
| Beetle | Cigarette | Foot | Kite | Pepper | Spoon | Watering can |
| Bell | Clip | Football | Ladder | Pineapple | Star | Watermelon |
| Belt | Clock | Fork | Lamp | Pitcher | Stethoscope | Well |
| Bench | Clown | Fox | Leaf | Pliers | Stool | Wheel |
| Bicycle | Coat | French-horn | Leg | Plug | Strawberry | Wheelchair |
| Bird | Comb | Frog | Lemon | Pot | Stroller | Whistle |
| Bone | Corn | Frying pan | Light bulb | Pumpkin | Suitcase | Window |
| Book | Couch | Funnel | Lion | Rabbit | Sun | Wineglass |
| Boot | Cow | Giraffe | Lips | Raccoon | Swan | Wrench |
| Bottle | Crab | Glass | Lock | Refrigerator | Sweater | Zebra |
| Bow | Crown | Glasses |  |  |  |  |

Naming was performed using simple line drawings of objects (derived from[36]). Item level analysis of these stimuli was performed in two primary ways. The first was done using measures of selectivity (number of possible correct responses) that have been characterized in normal subjects (derived from[36]). For the second, we used lexical frequency derived from a catalogue derived from natural speech[37]. We also sought to assess and control for additional variables including word length (# of phonemes), phonotactic frequency (average transition probability among adjacent phonemes in a word), and neighborhood density (# of phonemically similar words). Phonotactic frequency and neighborhood density values were calculated using the Irvine Phonotactic Online Dictionary[38].

During ECoG recordings, patients were able to verbally articulate their responses, but during fMRI acquisition patients were asked to internally (covertly) vocalize the word and respond with a button press that was recorded by the stimulus presentation software. For comparisons based on lexical frequency[37] and selectivity[36], object stimuli were ranked. Comparisons of high vs. low frequency were made using the 50 most frequent vs. 50 least frequent stimuli (in both the time-frequency and the functional connectivity analyses). A similar comparison was performed based on selectivity for each image.

### MR data acquisition

Imaging data acquisition was performed with a 3T whole-body MR scanner (Philips Medical Systems, Bothell, WA, USA) equipped with a 16-channel SENSE head coil. In all patients, the MRI data were acquired prior to surgery. Anatomical MR data were collected using a magnetization-prepared 180º radio-frequency pulses and rapid gradient-echo (MP-RAGE) sequence optimized for gray-white matter contrast with 1-mm-thick sagittal slices and an in-plane resolution of 0.938 x 0.938 mm. Functional MRI volumes (thirty-three axial slices, 3-mm slice thickness, 2.75 in-plane resolution, TE 30 ms, TR 2015 ms, flip angle 90°) were collected during performance of the language tasks, with stimuli arranged in a block design. Prior to data collection, each subject underwent pre-scan training using similar but non-identical stimuli for each task. Participants completed two runs of fMRI data collection. Each run was composed of eight blocks (136 volumes per run), and each block consisted of 10 task stimuli and 7 scrambled stimuli (20 s of task, and 14 s of scrambled control images). In total, 160 individual object naming stimuli and 112 scrambled stimuli were presented. Visual stimuli were presented at the onset of each functional image volume with Presentation software (version 11, Neurobehavioral Systems) using a screen positioned above the eyes (IFIS, Invivo; 1500 ms on-screen, 515 ms inter-stimulus interval, subtending a visual angle of ~10° x ~10°). Patient responses were monitored in real time using a fiber optic response pad connected to an interface unit (Current Designs), and by video monitoring of the patients face using a closed circuit television.

### Image analysis

Anatomical image realignment, spatial normalization transformation and group analysis were performed in AFNI[39]. Surface reconstructions of the gray/white matter interface and pial surface were generated using FreeSurfer v4.5[40]. fMRI volumes were aligned with a skull-stripped anatomical MRI. The aligned 4D dataset was spatially smoothed with a 3 mm Gaussian filter and then processed using multiple regression at each voxel to contrast the task (object naming) with the control condition (scrambled naming). To verify left-hemisphere language dominance, language lateralization indices were calculated for each individual using the language fMRI data[34]. Activations in Brodmann areas 44 and 45 in each hemisphere were extracted using masks constructed from a standard atlas[4]. The number of significant voxels (p<0.001, uncorrected) during the object naming was computed for the mask in each hemisphere. The laterality index used was equal to (#L - #R)/(#L + #R), where #L and #R are the numbers of significant voxels in the left and right hemispheres, respectively. A positive laterality index indicates left hemisphere dominance (Table 1).

### Subdural electrodes

Subjects were semi-chronically implanted with arrays of subdural platinum-iridium electrodes (PMT Corporation, Chanhassen, MN, USA) with a top-hat design (4.5 mm in diameter, 3-mm-diameter contact with cortex) embedded in silastic sheets (10-mm center-to-center spacing), using standard neurosurgical techniques[41]. After SDE implantation, an axial CT scan was acquired (0.5 x 0.5 mm in-plane resolution, 1-mm slices) and registered to the pre-operative, anatomical MRI. Each SDE was initially localized using the CT scan and then transformed to the MR imaging coordinate space[42].

SDEs were then localized on individualized, automatically parcellated cortical surfaces, and categorized based on the sub-region they overlay. Time-frequency decomposition of the ECoG signal was used to extract high gamma band power (70-110 Hz) at each electrode. High gamma band activity derived from ECoG data robustly tracks activity across cortical regions[26, 43] and strongly correlates with the BOLD signal[34]. Gamma band activity levels were averaged within each sub-region for each stimulus type to synthesize results across the group. Power change in the gamma band was then used to characterize responses in each sub-region to high vs. low semantic and selection loads, on a millisecond scale.

### Electro-corticography

ECoG recordings during language tasks were performed between three to seven days after grid implantation. Stimulus presentation was conducted using identical stimuli and the same Presentation software as used for fMRI data acquisition on a PC laptop. In all patients, large numbers of trials of object (~200) and scramble (~100) naming were performed. Stimuli were displayed at eye-level on a 15’’ LCD screen placed at 2 feet from the patient (2000 ms on screen, jittered 3000 ms inter-stimulus interval; 500×500 pixel image size, ~10.8° x 10.8° of visual angle, with a grid overlay on 1300×800 pixel white background, ~28.1° x 17.3° of visual angle). For each condition, images were randomly selected from our database and never repeated, so that each subject saw a unique sequence of images, with scrambled images randomly interweaved. Subjects were instructed to overtly name objects, and say, “scrambled” for the scrambled faces. A transistor-transistor logic pulse triggered by the Presentation software at stimulus onset was recorded as a separate input during the ECoG recording to time-lock all trials.

Audio recording of each ECoG session was used to accurately measure the onset of articulation and to compute reaction time. Initial estimates for the articulation onsets in the remaining trials were first determined using in-house software (written for MATLAB 2013a, The MathWorks, Inc, Natick, MA) to identify the first time point following stimulus onset in every trial at which the baseline amplitude was exceeded by 50% in each patient’s audio trace. Audio recordings were then systematically reviewed to precisely mark articulation onset times, which were then used to derive the reaction time (RT) for each trial. It is important to note that a natural occurrence in human subject data is the existence of variability both within and across individuals in RT. To minimize the effect of this confound, we first eliminated all trials with RT’s shorter than 600ms or longer than 2000ms. We performed all ECoG analyses twice, aligning data in two different ways: 1) aligning trials to stimulus onset, and then 2) aligning trials to articulation onset. In this fashion, we could ensure that neural processes engaged during both stages of analysis (i.e. following stimulus onset and leading up to articulation) were more likely to be similar across both epochs and individuals.

ECoG data were also visually inspected for inter-ictal epileptiform discharges and electrical noise. For 20 patients, ECoG data were collected at 1000 Hz during naming using NeuroFax software (Nihon Kohden, Tokyo, Japan; bandwidth 0.15-300 Hz). The other 7 patients underwent ECoG data collection at 2000 Hz (bandwidth 0.1-750 Hz) using the NeuroPort recording system (Blackrock Microsystems, Salt Lake City, UT, USA).

Electrodes were referenced to a common average of all electrodes except for those with 60-Hz noise or epileptiform activity when initially referenced to an artificial 0 V[44]. The data were imported into MATLAB, and the patients’ articulation times were extracted using the time-locked audio-video recording. To avoid including any brain regions with potentially abnormal physiology, all electrodes that showed inter-ictal activity (spikes) or that were involved with seizure onset were excluded from further analyses. All electrodes with greater than 10 dB of noise in the 60 Hz band were also excluded.

### Time series analysis

Spectral analysis was performed using the Hilbert transform and analytic amplitude to estimate power changes in different frequency bands (using MATLAB 2012b, Math Works, Natick, MA, USA). For time frequency analysis, the raw ECoG data were bandpass filtered (IIR Elliptical Filter, 30 dB sidelobe attenuation) into logarithmically spaced bands from 2 to 240 Hz. A Hilbert transform was applied, and the analytic amplitude was smoothed (Savitzky-Golay FIR, 2nd order, frame length of 128 ms) to derive the time course of power in each band. The percent change and t-value at each timepoint were calculated by comparing power to the pre-stimulus baseline (−700 to −200 ms). The percentages of change for all electrodes in each sub-region were then averaged together across subjects, and significance was assessed using a sign test and FDR corrected. Gamma band (70-110 Hz) activity was calculated using the same routines and compared between conditions (object vs. scramble, high vs. low frequency/selectivity) using paired t-tests.

### Functional connectivity

Functional connectivity was assessed using amplitude envelope correlations (AEC)[45-48]. Raw data were band-pass filtered in the frequency domain between 70 and 110 Hz using a square filter with sigmoid flanks (half amplitude roll off of 1.5 Hz) and were Hilbert transformed. An inverse Fourier transform was applied, and the absolute value smoothed with a moving average (100 ms long) to obtain the amplitude envelope of the signal. A noise correlation between pairs of channels was computed with the Pearson’s correlation at each time point across trials. To estimate the directionality of connectivity, the time series on one channel was lagged prior to AEC[49, 50] computation. This procedure could be performed in both directions (either channel lagged relative to the other) and at a variety of different latencies (from −250 to 250 ms in 10 ms steps). For individuals, significance was estimated using bootstrapping[50] (trial re-shuffling, 1000 resamples).

The gamma frequency range was chosen due to rapid signal attenuation (<6 mm) caused by high SDE impedance and low signal amplitude (2-5 µV for the 70-110 Hz band). The trial-by-trial fluctuations used to calculate AEC comprise a fraction of overall gamma power change and would presumably have an even greater fall off. This would, therefore, be even less likely to include signal overlap or dependence between channels. This high spatial resolution was evident in the presence of both negative and unidirectional correlations on adjacent channels. Such asymmetries strongly suggest that common sources are not an issue with these data or this analysis. Such assertions may not hold for lower frequencies because signal power is substantially higher and has greater spatial distribution. For these reasons, those frequencies were excluded from the current work.

### Grouped functional connectivity

In each individual, the SDEs localized in each area (POr, PTr, POp, sCG) were used to build a list of all possible pairs between any two sub-regions. Given that each patient may have multiple SDEs in an area, some subjects would have more pairs than others. For instance, if a subject had two SDEs in both POr and PTr, there would be four different combinations, whereas another subject with a solitary SDE in each of those cortical areas would only have one combination. It was necessary to limit the number of pairs from each person contributing to the group average to avoid having any given subject over-contribute to the group results. To do this, SDEs were only considered eligible if they met relatively low activation thresholds. For left PTr, POp, and sCG, a 15% increase in gamma power relative to baseline was considered ‘active’, whereas in left POr, SDEs were included if there was a 10% decrease. These criteria reduced the number of SDEs in each region to 22 POr sites, 22 PTr sites, 19 POp sites, and 28 sCG sites. The individual AEC results were computed for all SDEs in each individual, and they were then transformed into a Fisher’s z score, averaged and assigned significance[50]. For connectivity between each sub-region there were 28 POr/PTr pairs, 23 POr/POp pairs, 31 POr/sCG pairs, 29 PTr/POp pairs, 39 PTr/sCG pairs, and 41 POp/sCG pairs.

### State changes in the IFG and sCG during word generation

A K-means clustering of the AEC connectivity between all regions was performed [51, 52]. The goal of the clustering analysis was to incorporate correlations at positive lag, negative lag and 0 ms lag intervals and identify time windows (e.g. states) during which distinct inter-regional connectivity patterns can be identified (relative to subsequent or previous time windows). For this we chose three different time lags: −200, 0, and 200 ms. The −200 and 200 ms lags were chosen to minimize the amount of shared information they have with a lag of 0 ms. Lags for use in the cluster analysis were determined by cross correlating the AEC time series at different lags with 0 ms. We found that the correlational value decreased and plateaued around + or - 200 ms. Although only three lags were included (−200, 0, and 200ms), increasing the number lags to include more intermediary lags (e.g. 50, 100, 150ms) in the analysis did not change the results. Convergence and optimal cluster order were validated using silhouette plots, via the squared Euclidean norm as the distance metric, to control data over-fitting (kmeans and evalclusters function, MATLAB 2013b, The MathWorks) [52, 53].

Given four sub-regions, six different connections were possible (POr/PTr, POr/POp, POr/sCG, PTr/POp, PTr/sCG, and POp/sCG), and given that connectivity patterns for three different lags were considered (−200, 0 and 200 ms) - 18 AEC time courses were used for the analysis. Using K-means clustering, these time-series (−250 to 1250ms, in 10ms steps, after stimulus onset) were grouped temporally (151 data points across 18 dimensions) to find patterns of connectivity[52].

## Results

High-frequency (1-2 kHz) electro-corticographic (ECoG) recordings were collected in 27 patients with 3351 implanted subdural electrodes (SDEs). 210 of these SDEs were situated over the IFG (pars orbitalis - POr, Brodmann Area 47/10; pars triangularis - PTr, BA45; pars opercularis - POp, BA44) and the sub-central gyrus (sCG, BA6v/4) in both hemispheres. All patients performed the task within normal response parameters (Fig 2) and all possessed an average or higher IQ (mean IQ - 98±13). Only epochs corresponding to response parameters within normative ranges for both accuracy and latency (<2 s) were utilized.

**Fig 2.**
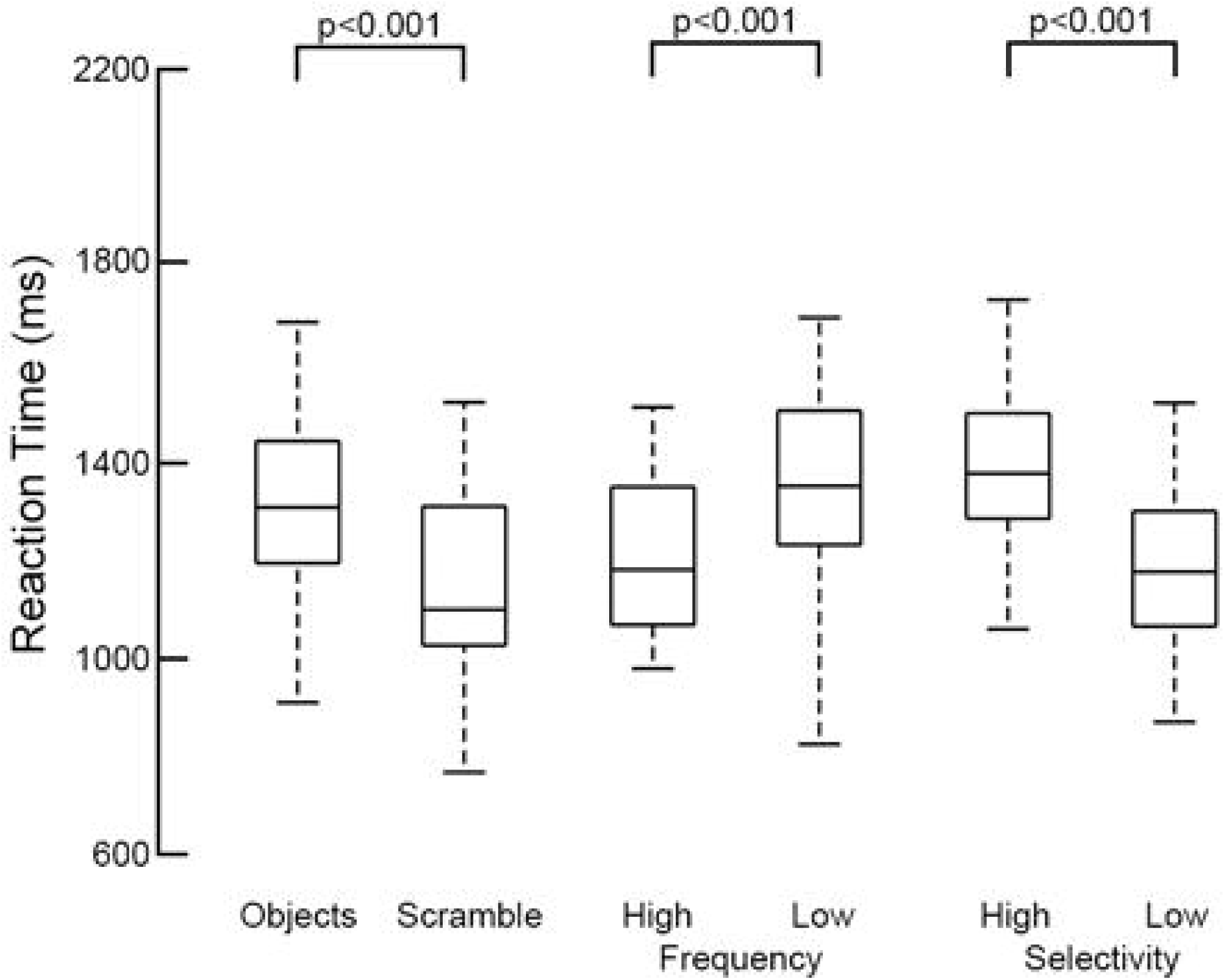
Reaction times per condition, across subjects (n=27). Using the audio recording, each patient’s reaction time was calculated based on the onset of their verbal response. Across the group, object naming had significantly higher latency than scrambled naming (p<0.001, two-sided, paired t-test. High-frequency objects had a significantly shorter latency than low-frequency words (p<0.001), and high-selectivity objects had higher latency than low-selectivity objects (p<0.001).

Event related spectral changes, particularly in the gamma frequency band were clearly seen in all sub-regions over the left hemisphere. For each individual and across the group, there was a marked decrease in gamma power in POr below baseline levels, starting 250 ms after stimulus onset and preceding activation seen in other sub-regions by ~100 ms (Figs 3 and 4; corrected p<0.01, two-tailed paired t-test). This decrease in gamma power was notable in both the object and scrambled naming conditions.

**Fig 3.**
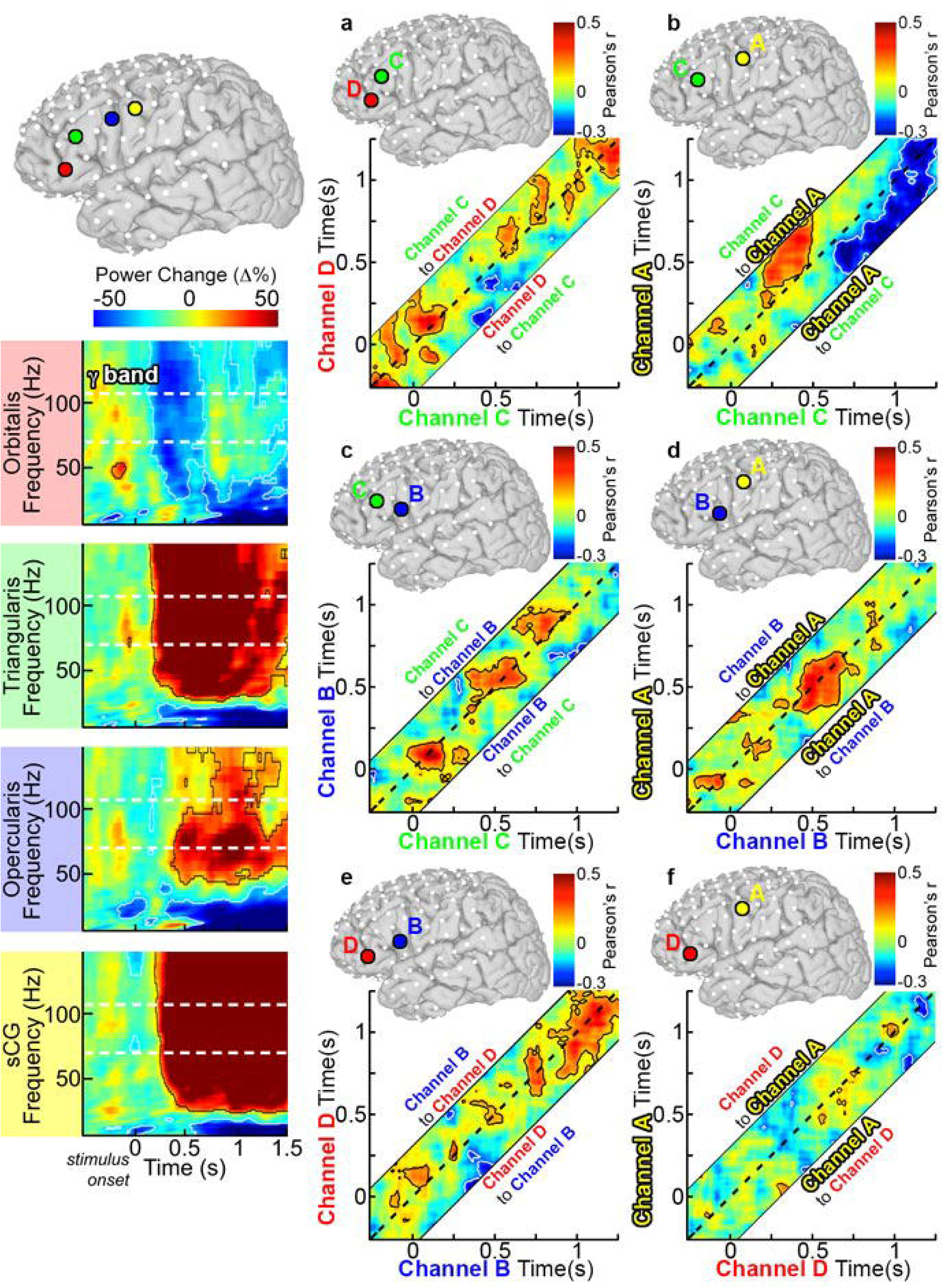
Single individual time-frequency graphs of activity during object naming. SDEs from a single, representative individual were anatomically localized to POr (red), PTr (green), POp (blue), and sCG (yellow). Time-frequency responses during object naming were computed and displayed percent power change relative to pre-stimulus baseline (two sided t-test, p<0.01 FDR corrected). Power changes at the individual level were consistent with those found across the group; including the POr gamma power deactivation and concurrent power increases in PTr, POp, and sCG. Using this same subject, connectivity analysis using the AEC method was also performed for these electrodes. (a) AEC was computed for two SDEs in a single subject over POr (Channel D) and PTr (Channel C). As in the group analysis, the dashed line represents a lag of 0 ms, areas above the dashed line represent activity on Channel C correlating with later activity on Channel A, and regions below the dashed line lag Channel D before Channel C. Confidence intervals were computed using trial reshuffling (contour lines are p<0.05 uncorrected, two-sided, 1000 resamples). (b) PTr and sCG, (c) PTr and POp, (d) POp and sCG, (e) POp and POr, and (f) POr and sCG.

**Fig 4.**
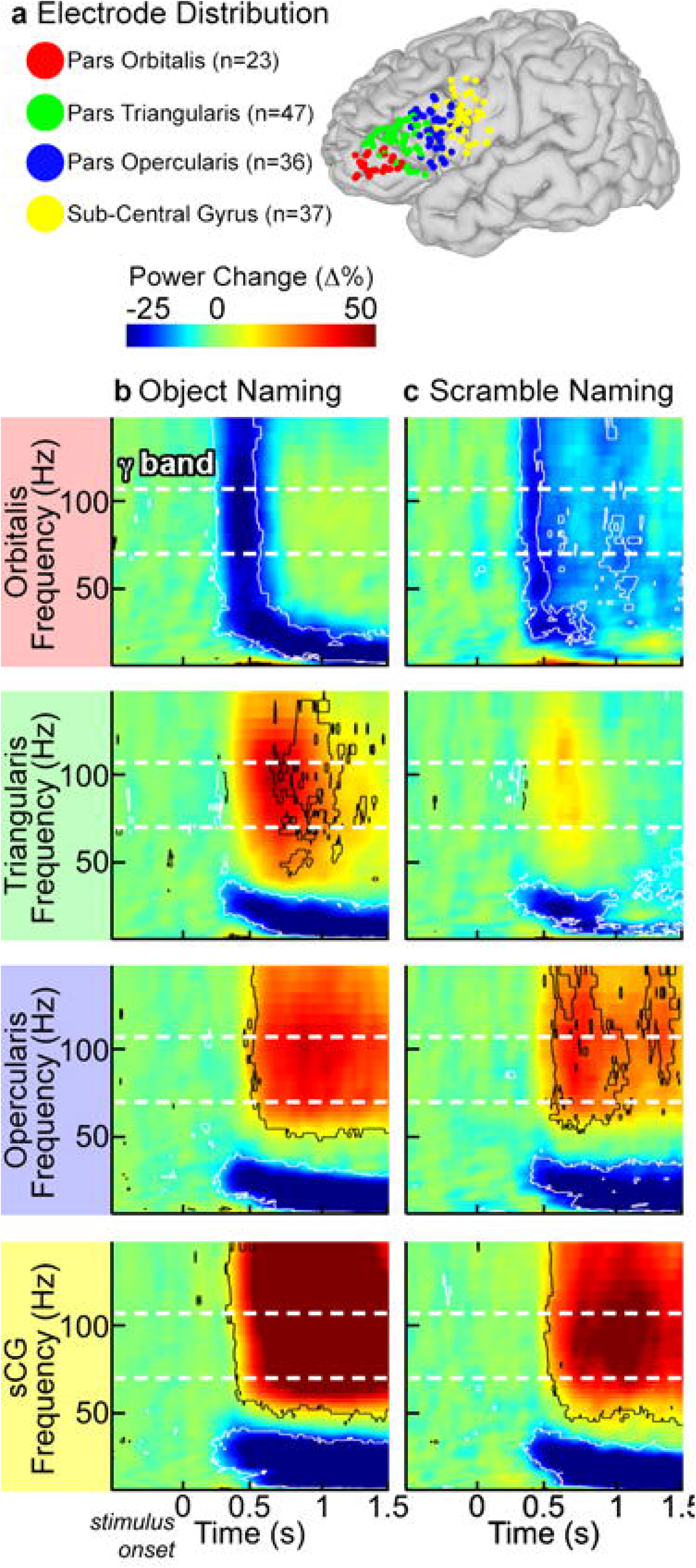
Time-frequency activity plots. (**a**) SDEs from 22 subjects with recording sites over the peri-sylvian cortex of the left hemisphere were anatomically localized to pars orbitalis (POr), pars triangularis (PTr), pars opercularis (POp), and sub-central gyrus (sCG), and then co-localized on a common brain surface. For both object (**b**) and scrambled image (**c**) naming, group time-frequency responses were computed by averaging the amplitude envelopes of gamma power for all SDEs across subjects in each sub-region (percent power change relative to pre-stimulus baseline, two-sided sign test, p<0.01 FDR corrected). During object naming, the first change from baseline was a decrease in high frequency power in POr, followed by concurrent increases in PTr, POp and sCG. The scrambled naming task showed weak activation of PTr, POp and sCG, although the time-points of these activations were similar for both tasks.

After this decrease, a prominent rise in gamma power occurred in both POr and PTr that was greater for object naming than for the scrambled image condition (Fig 5). A rise in gamma power was also observed in POp and sCG after the initial POr decrease; activity in these regions showed no effect of condition (object/scrambled) and varied only in the timing of the response. Gamma power increases in the anterior IFG (POr and PTr) were larger for object naming than for scrambled images, consistent with the notion that naming engages anterior IFG subregions more than simple articulation, perhaps due to semantic [12] or executive operations [54]. In contrast, the posterior IFG (POp and sCG) varied only minimally between naming and saying the word “scrambled”, consistent with the view that these regions code for lower-level phonological and motor processes.

**Fig 5.**
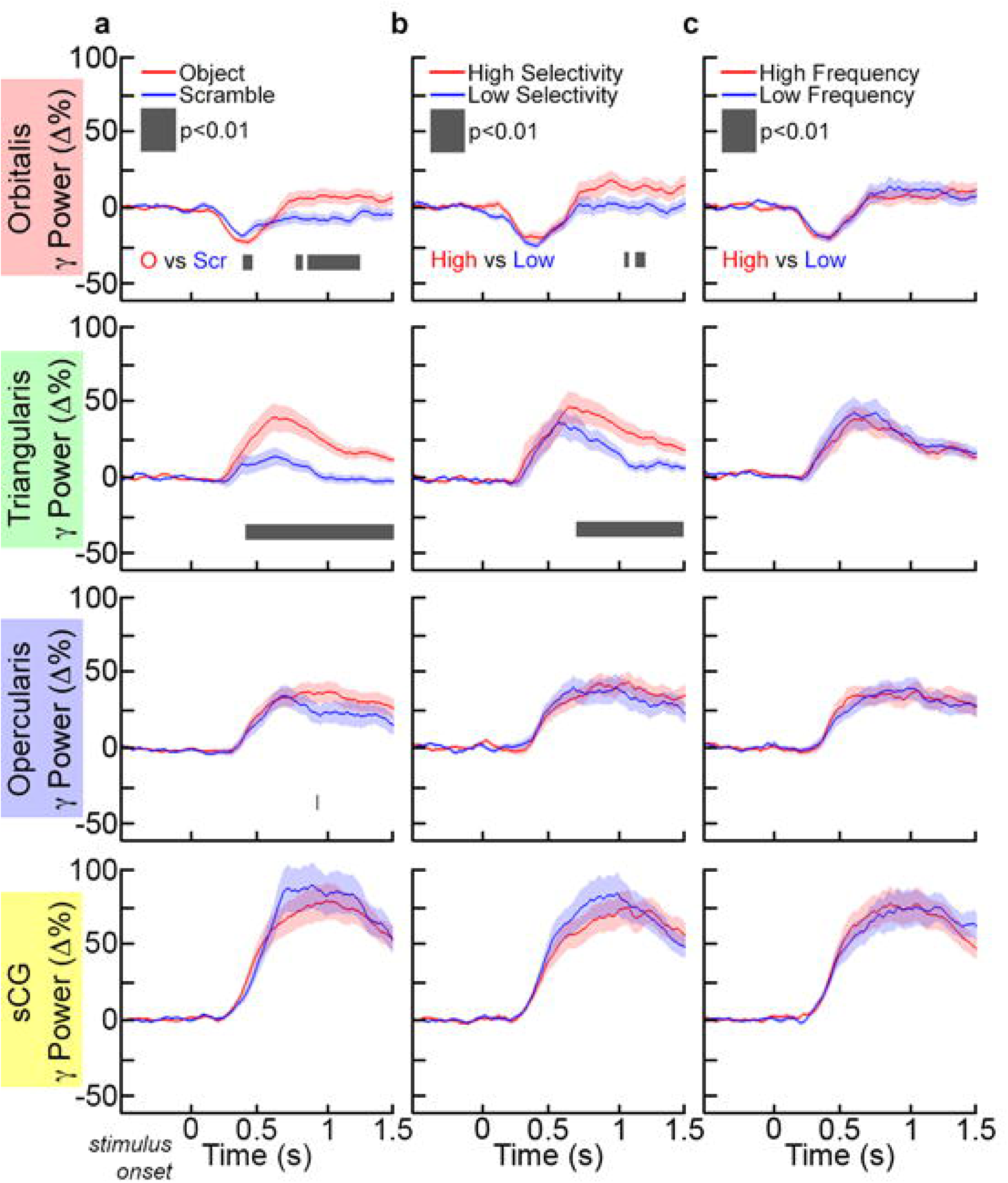
Gamma activation in IFG and sCG during each condition: Across subjects, gamma (70-110 Hz) power changes were averaged (mean±1SD) in each sub-region for (**a**) both tasks (object and scrambled naming). Comparisons between tasks and conditions were computed at each time-point (p<0.01, paired t-test, two-sided, FDR corrected). (**b**) In the comparison of high- or low-selectivity objects, only PTr showed a sustained difference in activation (black bar). However, (**c**) for high- or low-frequency objects, no regions showed a significant difference in activation.

The naming of stimuli with higher selectivity (greater number of possible correct responses) was associated with greater activation in PTr and POr relative to those with lower selectivity (significant at approximately 700 ms after stimulus onset), while no selectivity related differences were noted in either POp or sCG (Fig 5). The high versus low selectivity items did not differ in terms of phonotactic frequency (p = 0.51) or neighborhood density (p = 0.42). High and low selectivity items did differ, however, in terms of # of phonemes: on average low selectivity items had one fewer phoneme. We therefore assessed whether this variable contributed to the differences observed between high and low selectivity items by comparing gamma power time courses for long (6 or more phonemes) versus short (3 or fewer phonemes) items that were matched for selectivity. No significant differences were observed (Fig 6). Thus these data are concordant with functional imaging studies that localize lexical selection to the ventral IFG. Additionally, lexical frequency did not differentially modulate activity in any sub-region (Fig 5) of the IFG, also in agreement with prior non-invasive work [55].

**Fig 6.**
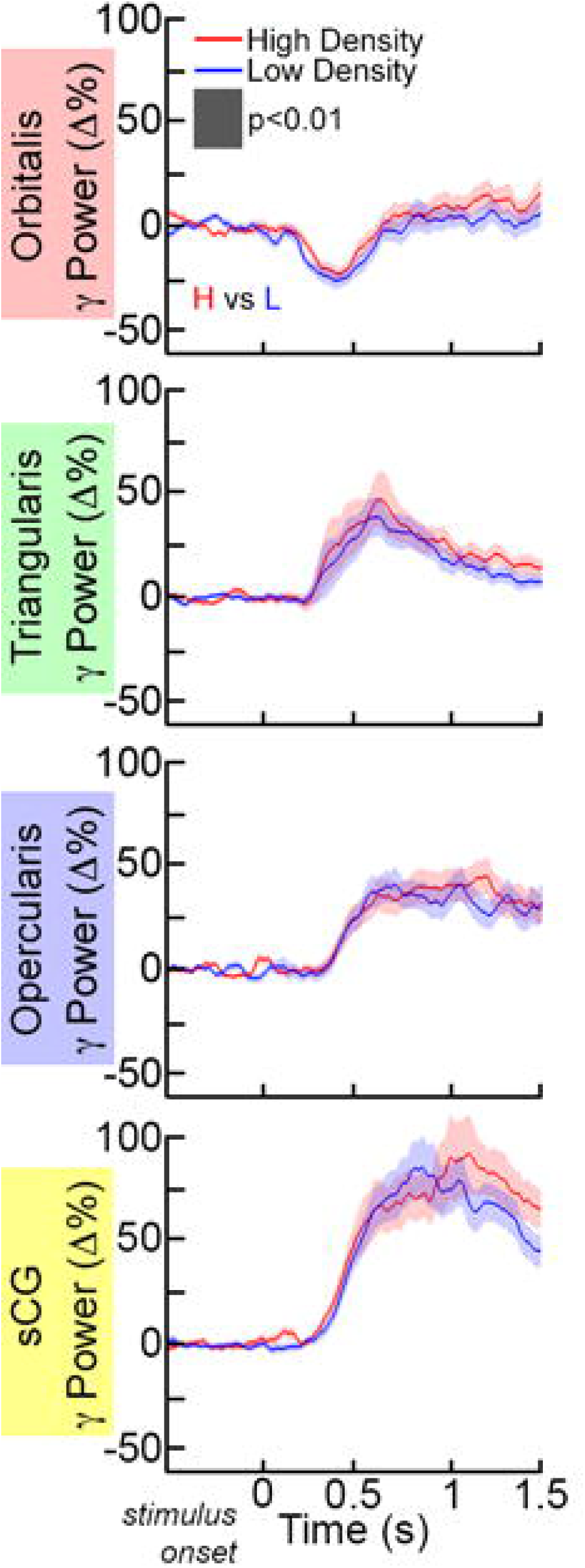
Gamma activation of IFG and sCG as a function of word length. Short items contained 3 or fewer phonemes and long items contained 6 or more phonemes. No significant differences were observed in Gamma (70-110 Hz) power changes in the group (mean±1SD) for short vs. long items (p<0.01, paired t-test, two-sided, FDR corrected).

An identical analysis using only data from non-dominant (in this case, all right sided) hemispheric recordings revealed that only the sCG was significantly active during these tasks (Fig 7), with minimal changes in the right IFG proper. The analysis did show some gamma activation in the right hemisphere PTr during the scrambled image condition, but this did not achieve significance. POr, PTr and POp in the language dominant (left) hemisphere were all significantly more active than their right hemispheric homologs (corrected p<0.05, two-sided unpaired t-test). The strongly lateralized nature of these electro-physiologic processes is consistent with the fact that all subjects were left-hemisphere-dominant for language.

**Fig 7.**
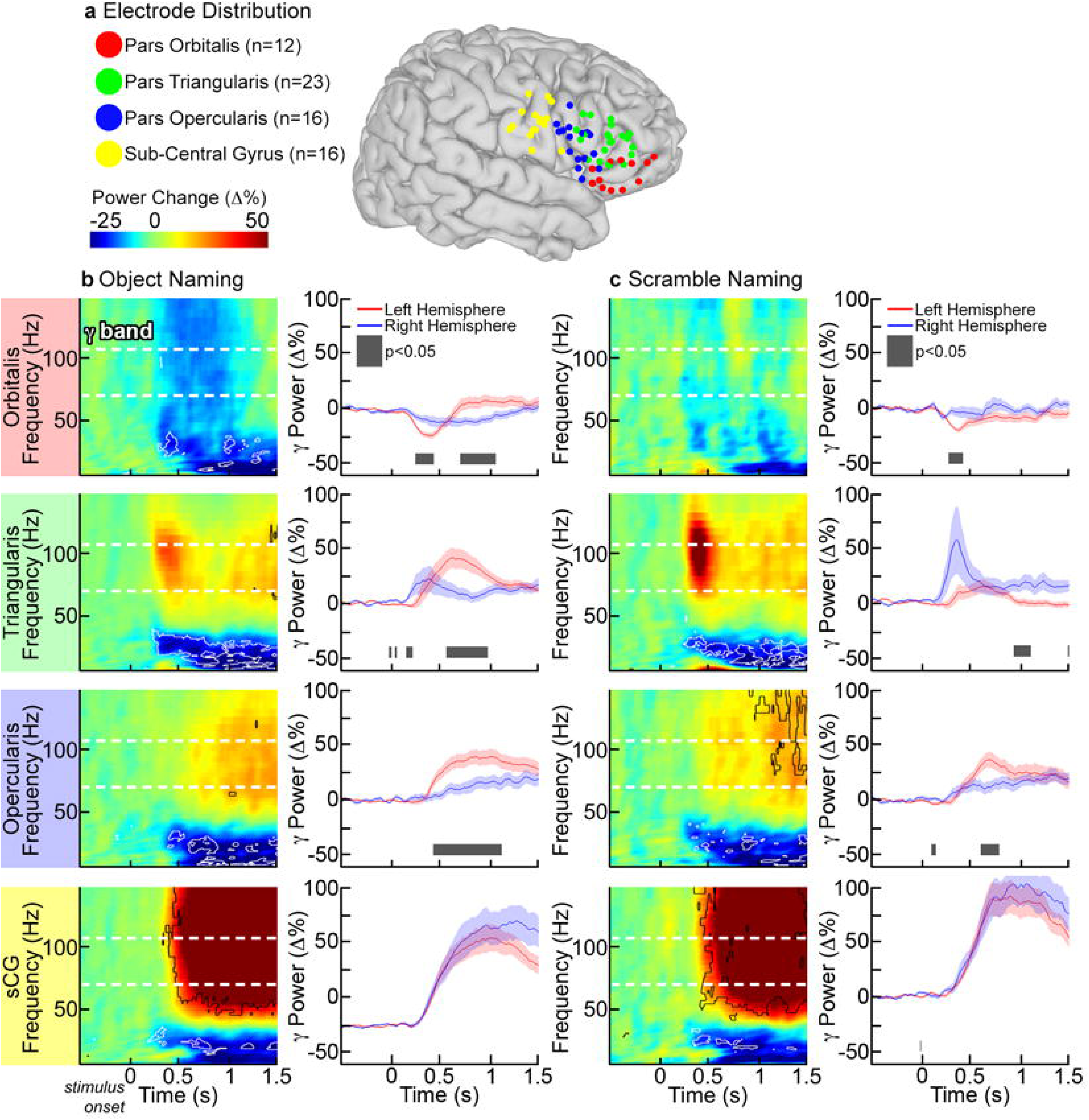
Time-frequency analysis of right hemisphere SDEs. (**a**) SDEs from 10 subjects with right-hemisphere coverage were co-localized on a common brain surface as shown previously (Fig 4). Grouped time frequency responses were computed for (**b**) object and (**c**) scramble naming (p<0.01 FDR corrected, two-sided sign test). A direct, left vs. right hemisphere comparison was made in the gamma (70-110 Hz) frequency range (p<0.05, unpaired t-test, FDR corrected) for all four regions. During object naming, POr, PTr, and POp were all significantly more active in the left hemisphere (black box). De-activation of left POr was also significantly different. Only two significant differences were noted during scramble naming: right PTr was more active in the comparison (but not against pre-stimulus baseline), and left POp was significantly more active.

An evaluation limited to the mean power reveals which regions are maximally engaged in which type of process, but provides minimal insight into the functional connectivity between these regions. To elaborate the brain dynamics governing rapid selection of appropriate verbal responses, we estimated coupling between sub-regions of the IFG. This was done using an amplitude envelope correlation (AEC) technique (Fig 1), chosen for its robustness to common source contamination, plus the modeling of connectivity at varied latencies[49]. Using lagged AEC in the gamma band power between active SDE pairs within each individual[45, 46, 49, 50] (significance tested by bootstrapping, 1000 resamples), we computed inter-regional functional covariance estimates for each subject, which we then averaged (after computing a z transform of individual r-values) across the group. If activation in one area correlated with and preceded activation of another area, this was taken to imply that the first region was functionally connected to the second.

At baseline (pre-stimulus rest), there was strong bidirectional connectivity observed between POr and PTr and between PTr and POp that continued till 300 ms after stimulus onset (Fig 8). However, resting correlations were weak between the other sub-regions, including pairs of regions that were immediately adjacent (e.g., POp and sCG), suggesting that this higher baseline connectivity of PTr with other regions was not simply a function of proximity but could indicate that this is an important computational hub[56].

**Fig 8.**
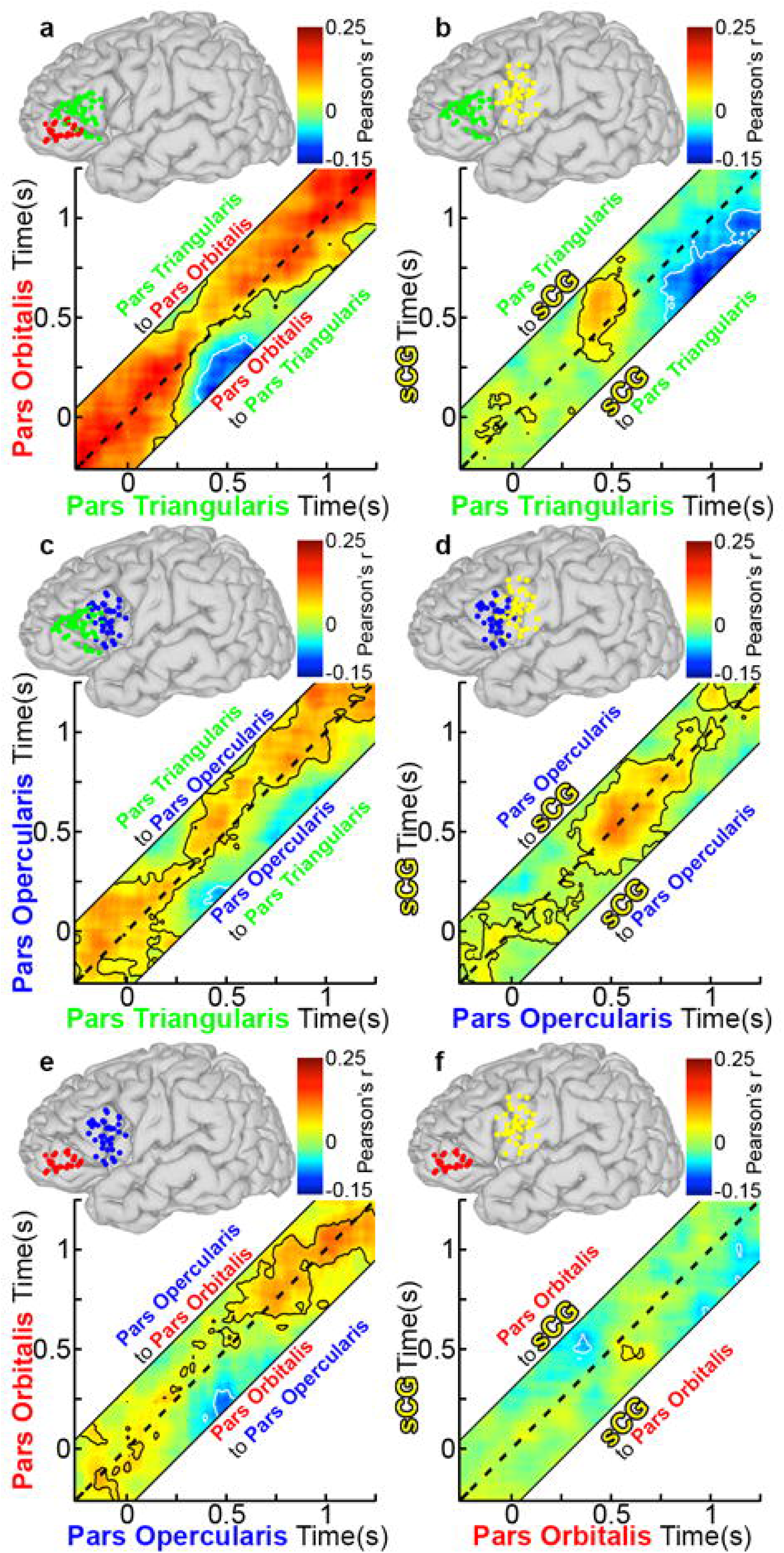
Functional connectivity of left IFG during object naming: (**a**) Grouped PTr and POr connectivity was estimated by averaging Amplitude Envelope Correlations (AEC) calculated for each individual and the averaged across the group (n=28 total pairs of SDEs, contour lines are p<0.05, two-sided t-test, FDR corrected). The dotted line on each graph represents a lag of 0 ms, while connectivity above or below this line is lagged from one region to the next (activity in region A affects region B at a later time point). See also methods and Fig 1 for further explanation. AEC across a range of lags (−250 to 250 ms, in 10-ms steps) are represented, with a dashed line depicting a lag of 0 ms. The area above the dashed line represents activity in PTr correlating with later activity in POr (directionality: PTr to POr), and the area below represents activity from POr correlating with later activity in PTr (directionality: POr to PTr). Warmer colors represent positive correlations between the two channels, and cooler colors indicate negative correlations. Unidirectional correlations appear as strong correlation either above or below the dotted line, while bidirectional correlations are centered on it. Given four sub-regions (POr, PTr, POp, sCG), there were six possible pairs of connectivity to evaluate: (**b**) PTr and sCG (n=39 pairs), (**c**) PTr and POp (n=29), (**d**) POp and sCG (n=41), (**e**) POp and POr (n=23), and (**f**) POr and sCG (n=31).

Between 300 and 600 ms after stimulus onset, lagged negative correlations (anti-correlated activity; p<0.01) were observed during object naming, in which activity in both PTr and POp was anti-correlated with activity in POr (Figs 8a and 8e). Similar but weaker correlations were observed during scrambled image naming (Figs 9a and 9e). Such a negative correlation between cortical regions could indicate either a direct increase in inhibition[8, 9, 57] or a dropout of excitation, resulting in a shift of the excitatory-inhibitory balance (a “de-excitation”). Baseline bidirectional connectivity between PTr and POp was reduced during this interval, and thereafter became principally unidirectional, from PTr to POp (Fig 8c). During the latter half of this window (between 400-600 ms) we also noted unidirectional, positive connections from PTr to sCG (Fig 8b). This interaction was followed by prominent, bi-directional coupling between sCG and the POp, which persisted through articulation (Fig 8d). No sharp temporal demarcation distinguished earlier POr-driven inhibition from later PTr-driven activation of the IFG. However, at 750 ms, just prior to articulation onset, sCG unidirectionally and negatively correlated with PTr (Fig 8b), an interaction suggestive of a signal to inhibit further lexical processes.

**Fig 9.**
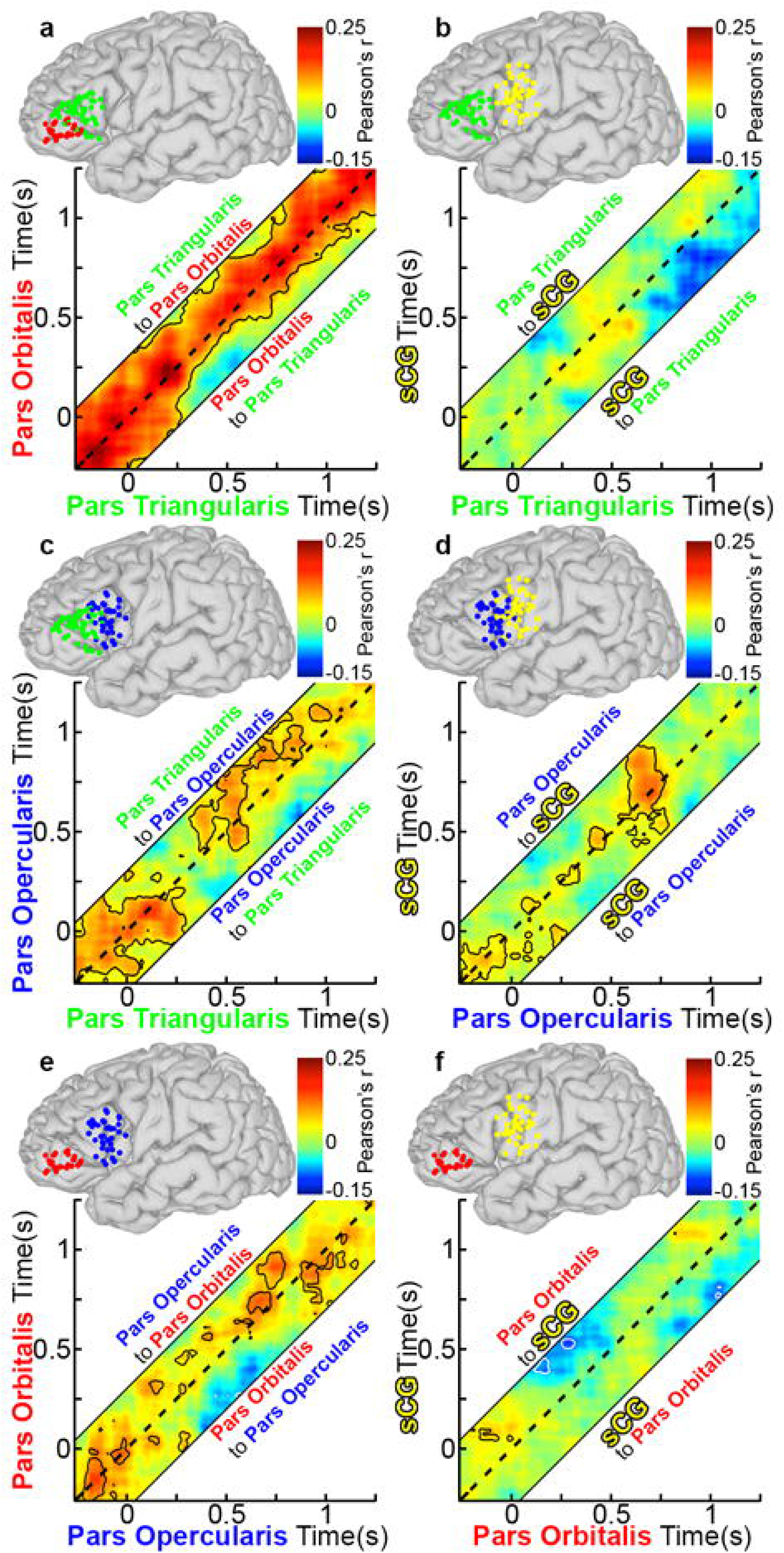
Functional connectivity of left IFG during scramble naming. Using the group AEC method in Fig 8, connectivity plots during scrambled imaging naming were made using the same electrode pairs as before. Overall, connectivity was weaker and shorter in duration during scrambled naming. (**a**) PTr and POr (n=28 total pairs of SDEs, p<0.05, two-sided t-test, FDR corrected), (**b**) PTr and sCG (n=39 pairs), (**c**) PTr and POp (n=29), (**d**) POp and sCG (n=41), (**e**) POp and POr (n=23), and (**f**) POr and sCG (n=31).

During the scrambled images condition, changes in correlations from the baseline state were weaker and transient (Fig 9). Specifically, both the significant inhibitory output from POr early in the process and the apparent stop signal to PTr from sCG at the onset of articulation were substantially weaker in this condition and did not reach significance.

The interactions described above are pair-wise. They reflect correlations between any two subregions at a time, but do not account for what other subregions are doing at the same time. In order to derive a holistic understanding of network behavior, beyond evaluating these individual pairwise inter-regional interactions, we integrated data from all interactions across the IFG during object naming. All pairwise amplitude envelope correlations at positive 200 ms lag, negative 200 ms lag and 0 ms lag intervals between all regions were evaluated concurrently to characterize network-wide transitions in patterns of connectivity (i.e. distinct states of inter-regional connectivity patterns) [51, 52] These 151 data points (analysis window: −250 to 1250ms, in 10ms steps, after stimulus onset) that lie in 18 dimensions (6 region-pair combinations, each with 3 distinct time-lag intervals), were reduced using k-means clustering[52]. We found that 4 clusters, representing distinct temporal states, were able to best characterize all 151 data points, and that there were smooth temporal transitions between these clusters. The four distinct temporal states are:

- A pre-stimulus to 300 ms post-stimulus stage with connectivity patterns similar to baseline
- An early stage (300-500 ms) during which the entire IFG becomes strongly interconnected, and which is highlighted by the negative correlation (possibly an inhibition) of PTr by POr,
- A late stage (500-750 ms) during which the posterior IFG and sCG become strongly coupled, and there is negative correlation (possibly an inhibition) of PTr by sCG
- Articulation-related activity (750-1250 ms) in sCG; connectivity within the IFG begins to return to baseline

The emergence of intermediate states in the functional connectivity of these regions suggests a novel conceptualization for the neural processes that occur during naming. Rather than a staged locus centric progression of activity from one region to the next, we observe distinct network states, i.e., distinct patterns of activation and connectivity between regions. Furthermore, these states may correspond with phenomenological ontology derived from older, behaviorally driven psycholinguistic models[17].

To depict both activation amplitude and connectivity during object naming, we represented gamma amplitude responses at each sub-region along with the inter-regional correlational activity for each state (each epoch derived from the K-means cluster), to generate a spatiotemporal map amalgamating activity plus connectivity within the system (Fig 10). This map demonstrates that inter-regional influences play critical roles in determining the extent and timing of gamma activity at each node in the system.

**Fig 10.**
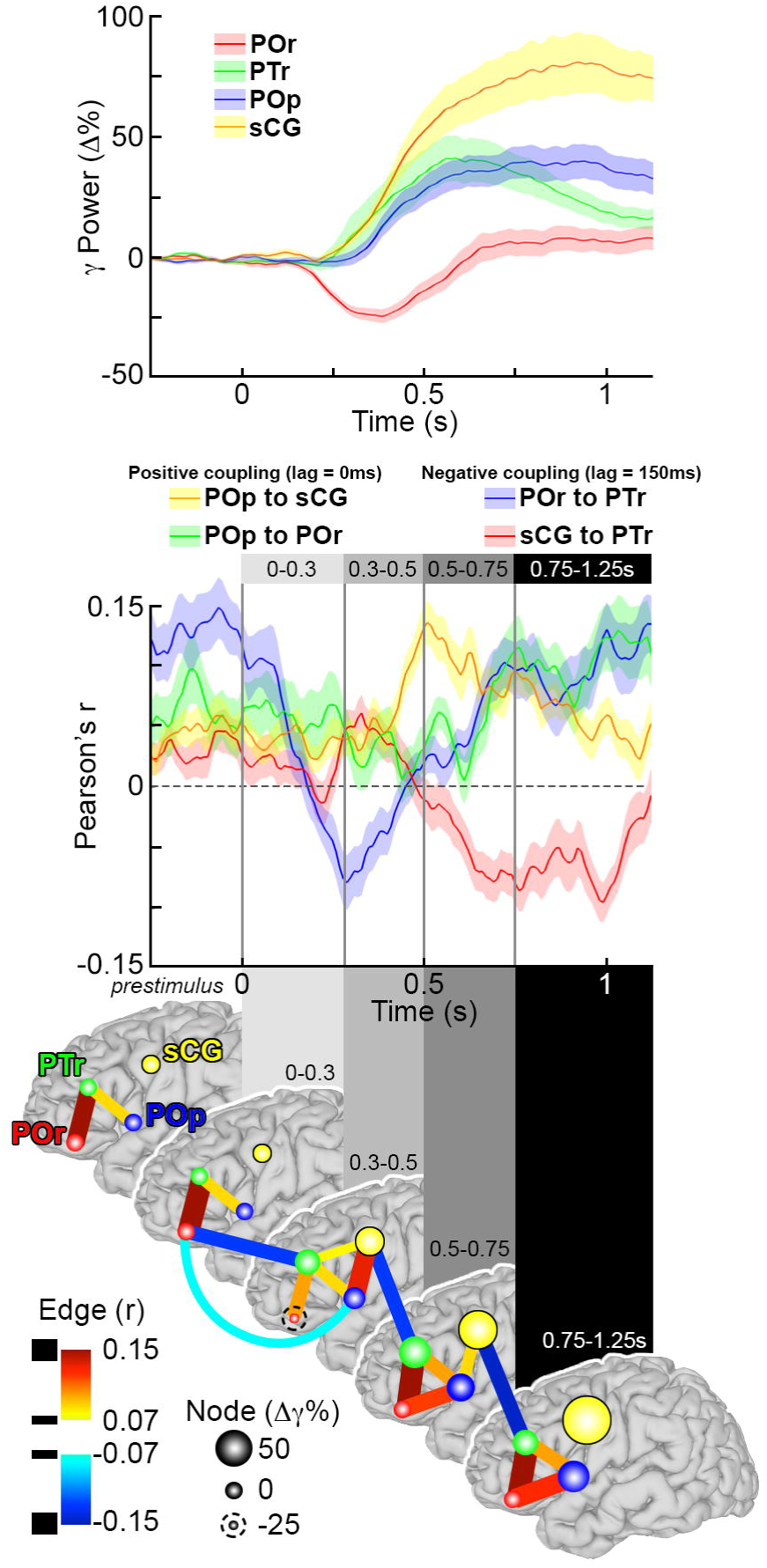
Co-representation of gamma power, connectivity and network states during word production. (**Top**) The first change in gamma power during object naming for all subregions in the left IFG show is a decrease in power in POr. This is followed 100 ms later by increases in power in PTr and sCG, and finally POp. (**Middle**) Four inter-areal interactions were selected to highlight state transitions in the IFG and sCG, two positive correlations: POp and sCG (lag=0 ms, Fig 8d) and POp to POr (lag=0 ms, Fig 8c), and two negative correlations: POr to PTr (lag=150 ms, Fig 8a) and sCG to PTr (lag=150 ms, Fig 8b). Each interaction represents one lagged time series of the AEC plot (mean across groups ± 1 SD). Time-points corresponding to network-state transitions were identified using a K-means clustering analysis, and distinct clusters are highlighted in different shades of gray. Following stimulus onset, four different time epochs were identified (0 to 300 ms, 300 to 500 ms, 500 to 750 ms, and 750 ms until vocal response). (**Bottom**) Using changes in gamma power during object naming and the network connectivity, a schematic of network dynamics was synthesized for object generation. These dynamics are shown at baseline and for the four subsequent processing stages, matching the network state transitions shown above. Each node represents the gamma power change for a given region, and the connections between them (edges) depict the inter-areal functional connectivity. Epochs are shown on separate brain surfaces, and connectivity within the epoch is at 0 ms lag. Connectivity that extends between adjoining surfaces is at non-zero (150 ms) lag.

Finally, given that these are all correlational measures and do not prove causality, we sought to validate the interpretations of inter-regional connectivity and the temporal boundaries provided by the k-means clustering during object naming. We therefore evaluated the influence of word frequency and selectivity on connectivity (Fig 11). We found that the negative correlation between sCG and PTr reached the same magnitude but occurred significantly earlier for high vs. low frequency objects (p<0.01, paired t-test). The onset of the negative correlation for each condition was within the late stage (500-750 ms). Given that reaction times were significantly shorter for high frequency objects, processing in the IFG likely terminates earlier for those stimuli. The same effect was observed for low-vs. high-selectivity objects (p<0.01, paired t-test). The finding that the negative covariance between sCG and PTr occurred earlier for words produced more quickly further supports the role of sCG in termination of processing within PTr[7, 58].

**Fig 11.**
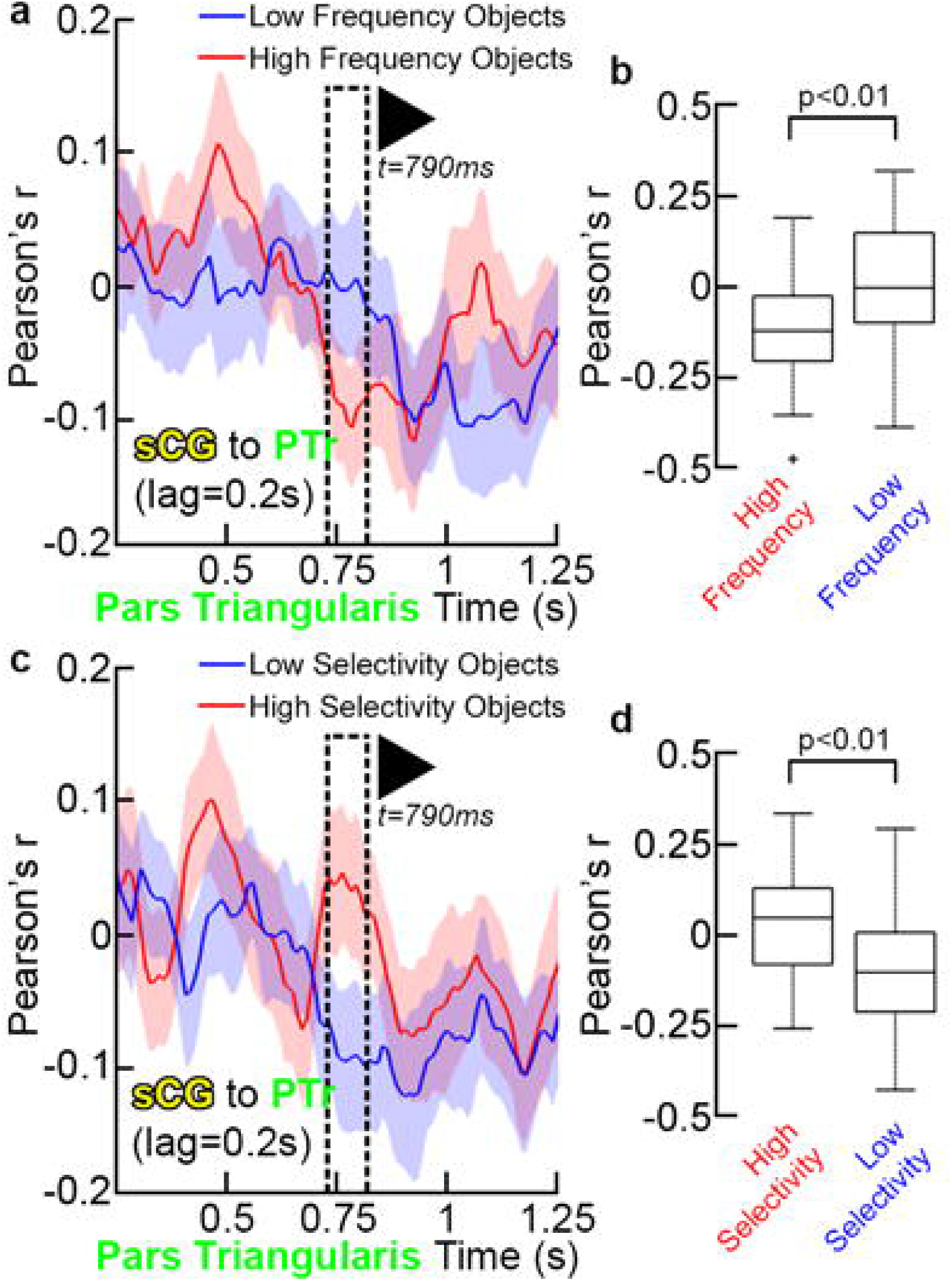
sCG interactions with PTr. (**a**) Functional connectivity between PTr and sCG were compared between high and low lexical frequency across the group (sCG to PTr, lag=200 ms, mean±2 SD, n=39 SDE pairs). Each line represents the correlation between sCG and PTr from pre-stimulus baseline through response onset at one lag. Just before articulation, this interaction becomes strongly negative; representing the feedback from sCG to PTr. Given the lagged timeline of correlations, the onset of sCG activity changes during this correlation occur 200ms prior to the time-scale for PTr shown here (**b**) When this correlation is compared for high frequency and low frequency objects, high frequency objects engage the feedback from sCG at a significantly earlier time-point (<800 ms) than low frequency objects (p<0.001, two-sided, paired t-test). This analysis was repeated for high- and low-selectivity objects. Given the lagged timeline of correlations, the onset of sCG activity changes during this correlation occur 200ms prior to the time-scale for PTr shown here (**c**) and (**d**), and demonstrated that the low-selectivity objects also engage the sCG feedback at a significantly earlier time-point.

Given that this stop signal occurs relative to the onset of articulation, we re-performed the time-frequency and functional connectivity analyses relative to vocalization. The initial vocal response for each trial was set at the time t = 0ms, and then trial were aligned at that point. The time-frequency responses were then averaged across individuals (Fig 12) locked to vocalization onset. We found that gamma power changes most closely associated with motor planning and control, e.g. those in POp and sCG, were most robust, while those in regions possibly involved in other process, POr and PTr, were much less significant than in the analysis time-locked to stimulus onset.

**Fig 12.**
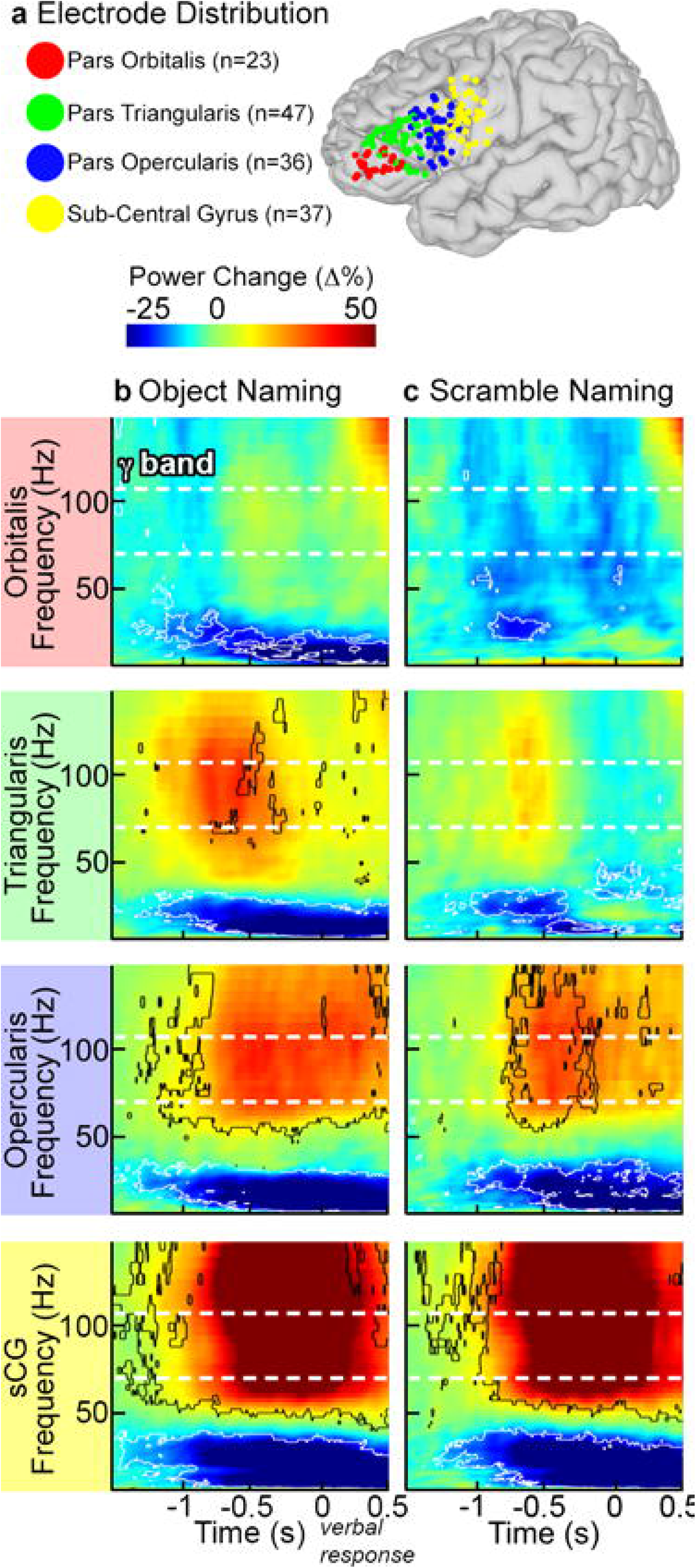
Time-frequency activity plots relative to articulation. (a) Using the SDEs located over POr, PTr, POp, and sCG, grouped time-frequency plots were recomputed during both object and scrambled naming relative to the onset of vocal response. In each individual, each trial was re-centered on the onset of articulation using the audio trace taken during the ECoG recording. The amplitude envelopes for all SDEs in each sub-region were then averaged across the group (percent power change relative to pre-stimulus baseline, p<0.01 FDR corrected, two-sided sign test). (b) In comparison with the stimulus onset locked analysis (Fig. 1), the most prominent gamma power changes occurred in sCG and POp – regions that are most closely associated with motor processing in speech production. Power changes in PTr and POr were diminished in amplitude. (c) During scrambled naming, only sCG and POp had significant power increases relative to baseline.

The analysis of connectivity using AEC method was also adjusted relative to articulation onset (Figs 13 and 14). The functional connectivity most associated with speech production - the negative correlation from sCG to PTr - was present here as well, beginning at 600 ms prior to vocalization (Fig 13b), persisting until articulation. This is consistent with the idea that it acts as signal to terminate PTr processing.

**Fig 13.**
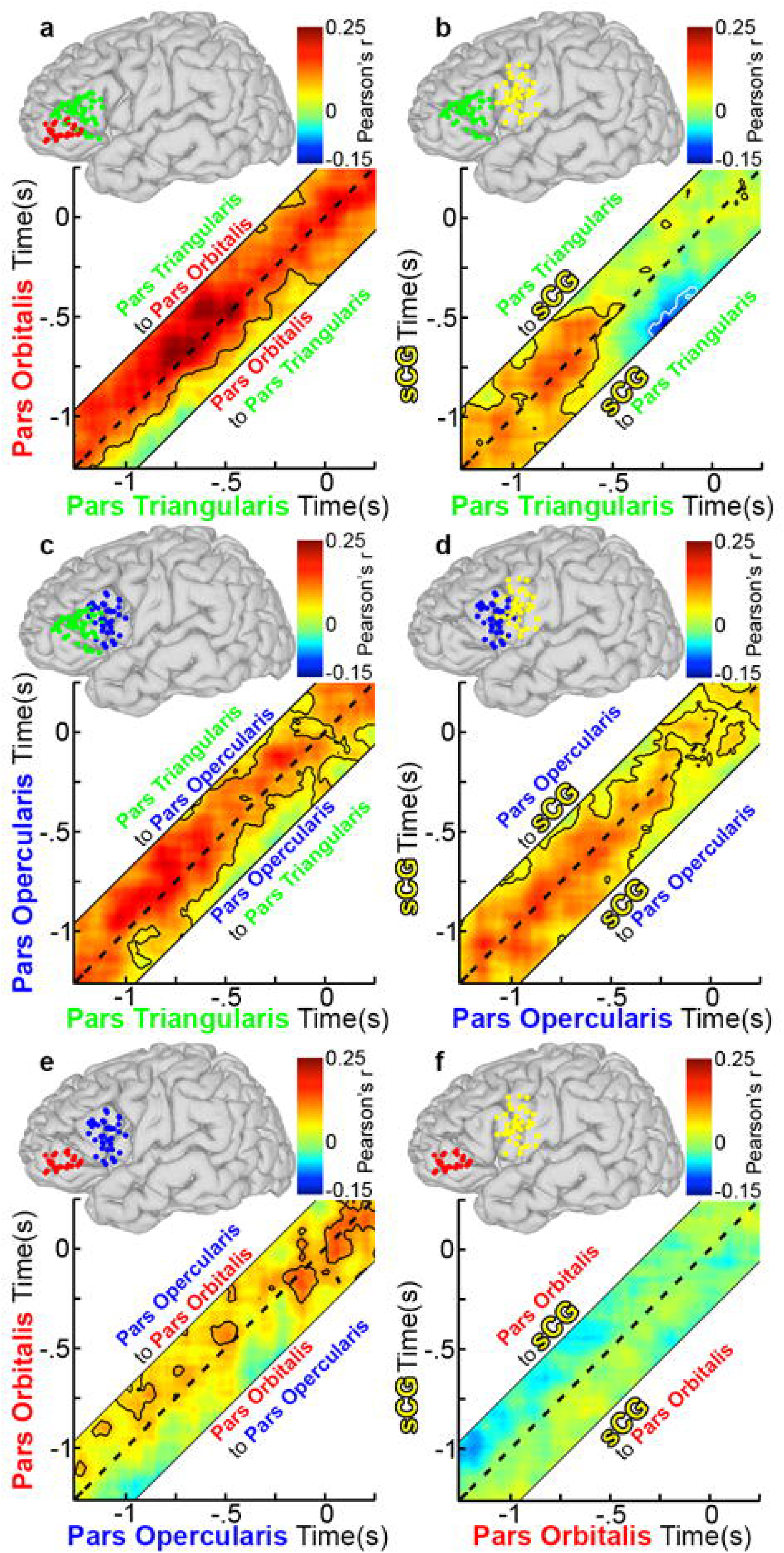
Functional connectivity during object naming relative to vocal response. Grouped connectivity analysis was recomputed relative to onset of articulation. This carried out by re-centering the data on an individual and trial-by-trial basis. From there, the group connectivity analysis was done as in Fig 3. (a) PTr and POr (n=28 total pairs of SDEs, p<0.05, two-sided t-test, FDR corrected). AEC across a range of lags (−250 to 250 ms, in 20-ms steps) are represented, with a dashed line depicting a lag of 0 ms. The area above the dashed line represents activity in PTr correlating with later activity in POr (directionality: PTr to POr), and the area below represents activity from POr correlating with later activity in PTr (directionality: POr to PTr). (b) PTr and sCG (n=39 pairs), (c) PTr and POp (n=29), (d) POp and sCG (n=41), (e) POp and POr (n=23), and (f) POr and sCG (n=31). Connectivity between POp and sCG, and PTr and POp were robust and bidirectional before articulation. The negative correlation from sCG and PTr was observed beginning around 500 ms before articulation.

**Fig 14.**
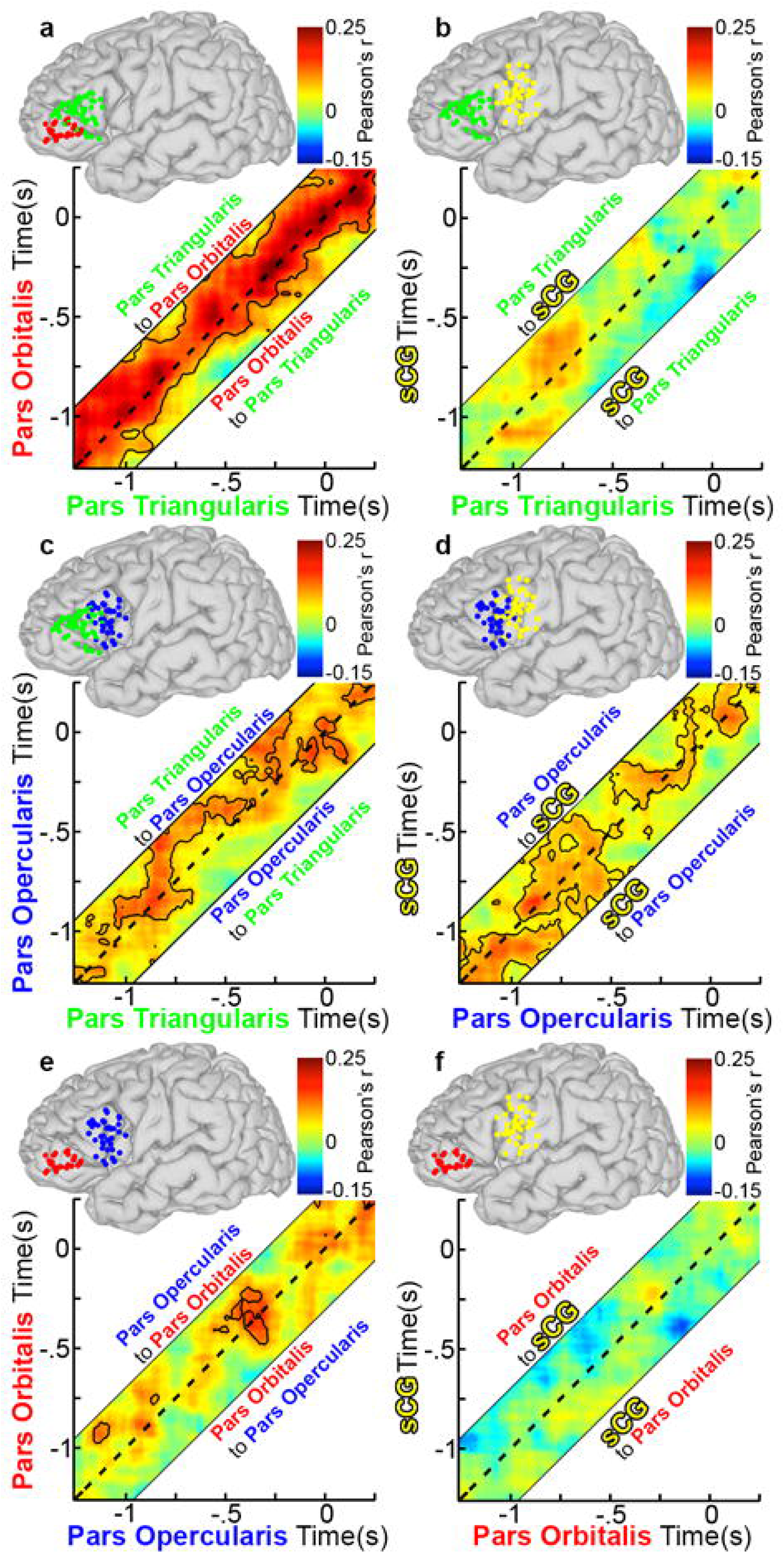
Functional connectivity during scramble naming relative to vocal response. Group connectivity analysis relative to onset of articulation was recomputed for the scrambled naming condition. (a) PTr and POr (n=28 total pairs of SDEs, p<0.05, two-sided t-test, FDR corrected), (b) PTr and sCG (n=39 pairs), (c) PTr and POp (n=29), (d) POp and sCG (n=41), (e) POp and POr (n=23), and (f) POr and sCG (n=31). Overall, magnitude of correlation was less also regions for the scrambled vs. the object naming conditions.

## Discussion

Research on the neuro-physiological basis of word production has been impeded by difficulties in capturing the temporal dynamics of rapid interactions between the distributed sites of language processing. In this study, we utilize high spatiotemporal resolution ECoG data, collected with dense intracranial recordings from a large cohort during object naming, to analyze patterns of network behavior in Broca’s area and surrounding regions of the language dominant frontal lobe. We find that that (1) cued word production entails a hierarchy of operations as suggested by psycholinguistic studies, but these are likely implemented by sequential states of a distributed network, not strictly serial processes in focal cortical regions[33]; (2) functional connectivity, rather than the magnitude of activity in individual sub-regions, provides greater insight into how speech is produced; (3) de-excitation and feedback-inhibition are the triggers for state-switching within IFG (between POr and PTr and between sCG and PTr); and (4) low level motor areas, such as sCG, are involved during the entire process from word selection to articulation.

The concurrent activation of PTr, sCG and POp seen here argues for a distributed, interactive parallel processing during word production, and against a serial unidirectional organization or a sequential activation process[59]. Cluster analysis of connectivity patterns further revealed distinct temporal states (i.e. clusters) during object naming (Fig 10). Based on pattern distinctions between conditions (object vs. scramble), as well as distinctions between high and low selectivity demands, we postulate that these states may reflect specific stages of word production. The pre-stimulus to 300 ms post-stimulus stage likely reflects a form of *attention modulation* - disengaging ongoing internally generated frontal lobe processes and facilitating the input of task relevant information. Visual and semantic processes in the ventral temporo-occipital cortex[44] are the likely drivers of this POr deactivation which then leads to the inhibition of PTr and POp via de-excitation. This process delays the onset of lexical retrieval presumably until processing in posterior regions of the cortex has matured. This view is concordant with prior theories of speech production both in that activation must be allowed to spread prior to selection, and that it is achieved through an interactive hierarchy between the visual processing and downstream control layers[29, 60].

Between 300-500 ms, activity and connectivity throughout the IFG and sCG was seen to increase. PTr is seen to unidirectionally engage sCG and POp, and bi-directional interactions between POp and sCG increase. These processes may represent the neural correlates of *lexical retrieval*. Subsequently, between 500-750 ms, *lexical selection and phonological code assembly* likely occur. POp/PTr are strongly functionally coupled and the posterior IFG and sCG become even more activated. sCG inhibits PTr during this interval, likely terminating lexical processing in PTr. Between 750-1250 ms, *articulation* begins. There is increased activity within sCG, while connectivity in the IFG starts to return to baseline.

The initial deactivation of POr seen here has also been shown by others to precede visual recognition[61]. This drop in high frequency oscillations in the IFG would be undetectable with techniques lacking sub-second resolution [62]. Given the invariance of this deactivation across conditions, it likely represents a conditionally independent modulation of ongoing prefrontal activity by inputs from visual and/or conceptual processing of the stimulus, and may allow for a state change from baseline activity to task relevant processing [61]. This deactivation leads to a broader modulation of activity in PTr and POp (Figs 8a and 8c). At rest, there is bi-directional positive connectivity between PTr and POr, but shortly after stimulus onset, excitation from POr to PTr selectively decreases, whereas inhibition does not. This de-excitation of PTr by POr likely enables processes outside the IFG to proceed uninterrupted[11]. These network interactions are consistent with a de-excitation model – a conjoined role of POr and PTr enabling *controlled retrieval*[8, 9], and may explain the inhibitory influence of POr on PTr in the context of decreasing POr activity[57].

In addition to response selection, there remains an unresolved debate regarding whether phonological processing is performed by PTr and POp together, or by POp alone (raised by theories regarding the relative roles of these regions [7] [31, 32]). PTr was significantly more active during object naming, whereas POp was equally active during both conditions (Fig 5). PTr also showed a “selectivity” effect, which ties it to a higher level of processing. The inter-regional interactions between PTr-POp-sCG were also weaker during scrambled naming (Fig 9). PTr was therefore less involved than POp in scrambled naming. Given the early activation in POp, it is clear that this region is more engaged in phonological operations than PTr. However, the early time-point at which POp is activated demonstrates that this process occurs in parallel with lexical retrieval and response selection.

Recent work suggests that Broca’s area becomes inactive at the onset of speech production; however, there was no description of the neurobiology that leads to this accomplishment[15]. Our analysis suggests that the underlying process relies on feedback from motor cortex to terminate further lexical processing. Although there were no differences in activation amplitude observed for high-vs. low-frequency object names, the functional coupling between sCG and PTr was significantly earlier for high-frequency stimuli. Similar patterns of results were observed for low and high-selectivity objects. That the negative correlation between sCG and PTr occurs earlier for high frequency stimuli corroborates the notion of stop signal from sCG to PTr[7]. The early involvement of sCG in word production and the feedback inhibition of PTr suggest that IFG interactions are more bi-directional than has previously been suggested.

We found that within the right IFG, no areas were significantly activated during word production (Fig 7). However, the right sCG did show significant increase in power similar to that found in left sCG suggesting a bilateral representation of the processes for articulation. The network interactions seen over the left hemisphere were also not present over the right side. The lateralization of activity in all patients to their language-dominant hemisphere (Table 1) implies that these are core language network interactions.

The map generated from this work provides a framework for lexical retrieval, selection, phonological code assembly, and articulation (Fig 10), but the map may be incomplete because as it does not incorporate visual and attentional processes or unsampled subcortical inputs to the articulatory system[63, 64]. Ultimately, all correlational methods for studying network behavior cannot prove causality or completely exclude the possibility of hidden common inputs. Additionally, in this work we did not consider long-range influences onto the VLPFC, both to limit computational complexity and due to varied sampling of distant cortical sites in these patients. Generating larger scale computational models of speech production will be the focus of future work.

It is important to note that our analyses were conducted using *noise* correlations, not direct power correlations. This type of correlation between signals recorded in different sites is specifically designed to minimize the possibility of type 1 errors when evaluating interactions between neural time series data. But it is important to emphasize that the effect sizes in noise correlations are substantially smaller than direct power correlations, and the range of the correlation coefficients we report here (between −0.5 and 0.5) remains consistent with an extensive and varied body of research in this field. Indeed, these studies have demonstrated that the reliability of noise correlations are best estimated by their significance as opposed to their absolute size. [65-72].

Our analyses were also performed time locked to stimulation presentation, but not to response vocalization, as the hypotheses to be addressed focus on response retrieval and selection, and not on articulatory mechanisms *per se*. Given that processes such as efference copy could be represented at multiple hierarchical stages and would likely vary with respect to stimulus features, they would likely confound analyses locked to articulatory onset[14]. Additionally, dissimilarities in initial syllables during response vocalization introduce marked variability in vocal onset times that we cannot currently account for without a large stimulus space (e.g. repeated trials per syllable type recorded for each patient). Finally, another source of uncontrolled variability is introduced due to differences in microprocessor-induced audio delays (5 – 50 msec range) inherent to the clinical recording system (Nihon Kohden) that was used in roughly half of our patient cohort. Although grouped ECoG time series analyses are robust to such variability, correlational analyses linked to articulation onsets are less reliable than those locked to stimulus onset. Future studies will address these issues.

Overall, our data reveal distinct hierarchical stages that have been postulated to occur during cognitive operations[60, 73], and add to a growing body of work supporting the parallelized and distributed behavior of the cortical networks underpinning speech production[27]. The time courses of activation in the individual IFG sub-regions show that power increases steadily over time in the lower-level areas (POp and sCG), ramps up then down in the intermediate level area (PTr), and has a complex biphasic profile in the highest (POr). However, we do not find any strictly serial activation profiles, or rigid feed-forward-only interactions in the prefrontal cortex[59, 74]. The observed interactions between IFG sub-regions change rapidly in magnitude, direction, and valence. Crucially, the functional status of a region is best described by inter-regional interactions, and not by the absolute level of activation in a given region (Fig 10). The framework of prefrontal – motor interactions during speech production and the de-excitation model outlined here may have broader implications for the study of pre-frontal control.

## Acknowledgments

We thank Adam Aron, Cathy Price, and Xaq Pitkow for comments on the manuscript, and Kenny Vaden for his assistance with the phonotactic and neighborhood density calculations. We are particularly indebted to the patients who participated in this study, the neurologists (Drs. Slater, Kalamangalam, Hope, and Thomas) who provide care to these patients, and the nurses and technicians in the Epilepsy Monitoring Unit at Memorial Hermann Hospital who helped make this research possible.

